# The unraveling of balanced complexes in metabolic networks

**DOI:** 10.1101/2021.03.23.436554

**Authors:** Damoun Langary, Anika Küken, Zoran Nikoloski

## Abstract

Balanced complexes in biochemical networks are at core of several theoretical and computational approaches that make statements about the properties of the steady states supported by the network. Recent computational approaches have employed balanced complexes to reduce metabolic networks, while ensuring preservation of particular steady-state properties; however, the underlying factors leading to the formation of balanced complexes have not been studied, yet. Here, we present a number of factorizations providing insights in mechanisms that lead to the origins of the corresponding balanced complexes. The proposed factorizations enable us to categorize balanced complexes into four distinct classes, each with specific origins and characteristics. They also provide the means to efficiently determine if a balanced complex in large-scale networks belongs to a particular class from the categorization. The results are obtained under very general conditions and irrespective of the network kinetics, rendering them broadly applicable across variety of network models. Application of the categorization shows that all classes of balanced complexes are present in large-scale metabolic models across all kingdoms of life, therefore paving the way to study their relevance with respect to different properties of steady states supported by these networks.

**Highlights:**

- Balanced complexes are ubiquitous in real-world networks, and facilitate insights in steady state flux phenotypes in large-scale metabolic networks.
- Novel factorizations are proposed that explain the formation of balanced complexes and enable their categorization.
- The results also provide a computationally-efficient tool for the identification of balanced complexes in large-scale networks.
- Examination of metabolic network models across all kingdoms of life shows that all categories naturally arise in this type of networks.

## 1 Introduction

The last two decades have witnessed the generation of large-scale metabolic networks along with the development of computational approaches within the constraint-based modeling framework that facilitate insights in the genotype – phenotype map (Bordbar, et al., 2014). Metabolic networks include the entirety of biochemical reactions through which multiple species (e.g. metabolites, metabolite-enzyme complexes) are taken up from the environment and/or are transformed into the building blocks of biological systems. Since the first stoichiometric models of *E. coli* (Varma & Palsson, 1994; Pramanik & Keasling, 1997), more refined metabolic models have been generated not only by increasing the gene and reaction coverage, but also by considering interactions of multiple cell types, tissues, organs in an organism (Martins Conde, et al., 2016) as well as interactions between organisms in communities (Fang, et al., 2020).

The success of the constraint-based modeling framework can in part be ascribed to: (i) invoking the simplifying assumption of steady state for the concentration of species and (ii) taking a flux-centered view that simplifies the metabolic constraints to a system of linear equations, in terms of the reaction rates (i.e. fluxes). The system of linear equation can be readily derived solely based on the stoichiometry of the analyzed network. Together with the capacity to include natural constraints on fluxes, these simplifications allow prediction of metabolic phenotypes with an assumed objective by employing classical convex optimization techniques (e.g. as done in flux balanced analysis (Orth, et al., 2010)).

The same quest for linearity is followed in chemical reaction network theory (CRNT) (Gunawardena, 2003). Like constraint-based modeling, CRNT also invokes the steady-state constraints; however, CRNT takes a concentration-centered view and often makes the simplifying assumption that the modeled reactions follow the classical mass action kinetics (Feinberg, 2019). As a result, one obtains a system of polynomial equations in terms of the species’ concentrations which can be analyzed at different levels of abstraction. For instance, linearity in CRNT is achieved due to the consideration of complexes, corresponding to the substrate / product side of each modelled reaction. This allows the rewriting of the systems of polynomial equation into linear equations in terms of the monomials, arising from mass action kinetics, parameterized by the rate constants of the assumed mass action kinetics.

As a consequence, CRNT has relied on linear algebraic tools to address very general questions about whether or not a dynamical property arises given any or some choices for the value of parameters (Feinberg, 1987; 1988). For instance, one of the seminal results in CRNT deals with so-called complex balanced networks in which every complex is balanced, in the sense that the sums of reaction rates using the complex as a substrate or product are the same at every steady state supported by the network. Complex balanced networks have been shown not to exhibit exotic dynamic behavior, including capacity for multistationarity (Feinberg, 1987) or presence of species showing robust concentrations (Shinar & Feinberg, 2011). Determining if a network is complex balanced can be done by calculating the deficiency of the network, relying solely on the structure of a given network (Feinberg, 1987; Gunawardena, 2003).

However, balanced complexes are not only present in complex balanced networks. For instance, under the steady-state assumption, a complex including a species that does not occur in any other complex of a given network is balanced (Küken, et al., 2021; Feliu & Wiuf, 2013). Therefore, it is natural to ask the following three questions: Can balanced complexes be efficiently computed in networks of arbitrary kinetics? What are the effects that realistic flux constraints have on presence of balanced complexes? What are the network mechanisms that lead to the presence of balanced complexes with and without invoking flux constraints? Answers to these questions go in the direction of reconciling the flux-and concentration-entered views, typical for the constraint-based and CRNT approaches. They also can help in shedding light on recent developments based on constraint-based linear programming approaches to identify balanced complexes in networks with arbitrary as well as mass action kinetics (Küken, et al., 2021). Since it has also been shown that balanced complexes can be used for effective reduction of large-scale metabolic models, answers to the abovementioned questions can help in addressing this problem that has gained some renewed interest (Singh & Lercher, 2020).

Here, we consider networks with arbitrary kinetics and provide a categorization of balanced complexes. The categorization delineates the contribution of network structure and flux constraints in categorizing a balanced complex into one of the four classes. We also formulate feasibility problems whose solutions can be used to categorize a balanced complex in a respective class. Our results also provide conditions that preclude the presence of particular class of balanced complexes. In addition, by analyzing high-quality metabolic networks across kingdoms of life we show that all classes of balanced complexes are prevalent and, therefore, relevant for properties of the supported steady states.

## 2 Material and methods

### Preliminaries on Chemical Reaction Network Theory (CRNT)

In the classical CRNT literature (Aris & Mah, 1963; Horn & Jackson, 1972; Feinberg, 1979; 1980; 1987; 1988), a chemical reaction network (CRN) is usually defined by the 3-tuple 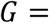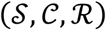, where 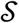 is a set of *m species/metabolites* and 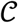 is a set of *n complexes*, whose elements 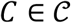 can be regarded as multisets of species, 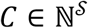. The set 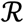 is formally defined as a relation between network complexes, 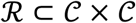; each element of 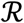 symbolizes the conversion of one complex to another and is referred to as a *reaction* of the network.

Throughout this manuscript, the standard basis for ℝ^*n*^ will be denoted {**e**_1_, **e**_2_, …,  **e**_n_}, where

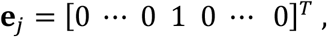

is an *n*-vector with a unit value at the *j*^th^ entry and zero entries elsewhere. Assuming some arbitrary ordering on the sets of species (*S*_1_, …,  *S*_*m*_), complexes (*C*_1_, …,  *C_n_*), and reactions (*R*_1_, … *R_r_*), any complex 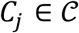 can be alternatively represented by a vector **e**_*j*_ ∈ ℝ^*n*^ indexing its position in the ordered set, and at the same time associated with a unique vector *y_j_* ∈ ℝ^*m*^, representing its species content. This defines the following mapping

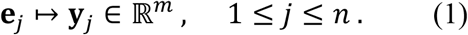

As a result, any given CRN is associated with a matrix defined as a compilation of vectors **Y** = [**y**_1_… **y**_*n*_], referred to as the *stoichiometric map*. In a similar fashion, one can associate any reaction 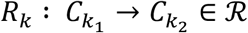 with vectors 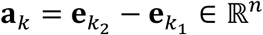 and 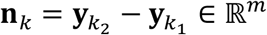. Assembling these vectors in the same order gives the *incidence matrix* **A** = [**a**_1_… **a**_*r*_] and the *stoichiometry matrix* **N** = [**n**_1_… **n**_*r*_], respectively. If follows that **N** = **Y A**.

The dynamics of the CRN is formulated by the following equation

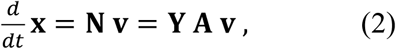

where 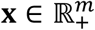 is the vector of species concentrations and **v** = **v**(**x**) ∈ ℝ^*r*^ is the vector of kinetic rates, referred to as *fluxes*, which is generally a nonlinear function of **x**. The *flux distribution* **v** is often bounded by an upper bound **v**_u_ and a lower bound **v**_l_ as follows

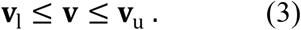

By convention, **v**_u_ has strictly positive entries for all reactions. The set of *irreversible reactions* 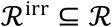 can be determined by the nonnegative elements of **v**_l_, that is

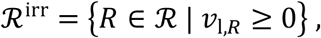

where 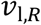 is the designated lower bound on flux through reaction *R*.

We say the system is operating in a *canonical flux regime*, if 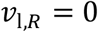, 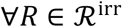 The set of *reversible reactions* is simply defined as the complement of 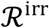 in 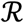, i.e. 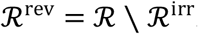. The system is said to be operating in an unbounded flux regime, if 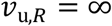, 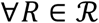 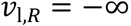, 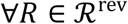. One can always choose an ordering of the reactions in which elements of 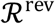 precede those of 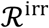. In this case, the incidence matrix **A** can be partitioned into matrix blocks **A** = [**A**^rev^ **A**^irr^].

For the system to be at steady state, the flux vector **v** must lie in the nullspace of matrix **N**, that is

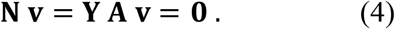

The intercept of the inequality constraints in Eq. (3) and equality constraint in Eq. (4) defines a convex set of feasible vectors **v**, *the steady state flux set*, denoted 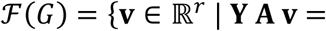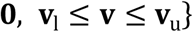. In a canonical and unbounded flux regime, the feasible set forms a polyhedral cone, referred to as the *steady state flux cone*.

For any 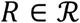, we say *R* is a *blocked reaction* in *G*, if for all the feasible set distributions 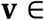 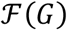, there is zero flux through *R*, that is, 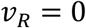 This is a commonly occurring phenomenon in metabolic networks, especially in scenarios when flux bounds and/or optimization of particular objectives are imposed. Similarly, we say a reaction 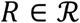 is *fixated* at some flux value *f*, if for all the flux distributions 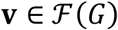, the flux through *R* is unchanged, namely, 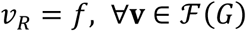. Clearly, any blocked reaction is one fixated at zero.

### 2.2 Balanced complexes

An important class of steady state fluxes are the so-called *complex-balanced steady states*, which were introduced by Horn and Jackson (1972) (Horn, 1972) as a generalization of detailed balanced steady states. At a complex-balanced steady state, the flux distribution satisfies

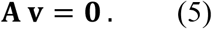

Eq. (5) constrains **v** to lie in ker **A**, which is generally a subset of ker **N**, and, hence, is a stronger condition compared to Eq. (4). From a conceptual viewpoint, complex-balanced steady states are those in which the algebraic sum of the total flux entering and leaving each complex equals zero. In more technical terms, the flux vector is composed of a number of cyclic generators (Johnston, 2014), i.e. nonstoichiometric elementary flux modes (Conradi, et al., 2007). Of particular interest are systems for which all steady states are complex-balanced, that is, Eq. (4) can be seamlessly replaced by Eq. (5). This class of systems -referred to as *complex-balanced systems*- are well-studied in the classical body of work on CRNT, and very strong results have been derived for them, most notably the well-known Deficiency Zero Theorem (Feinberg, 1987).

Unfortunately, in most cases those strong results cannot be applied to the majority of real-world metabolic networks: Due to the more complex nature of such systems, often not all but only some complexes are balanced at steady state. However, recent studies have shown that, even in absence of full complex-balancing, existence of balanced complexes can be exploited to obtain significant network reductions and throw some light on the steady states (Feliu & Wiuf, 2013; van der Schaft, et al., 2015; Küken, et al., 2021). This gives rise to the question regarding the underlying mechanisms that lead to the formation of complex balancing in metabolic networks.

Let us next give a formal definition of a balanced complex. For a given matrix **X**, let **X**^:*i*^ and **X**^j:^ denote the *i*^**th**^ column and j^th^ row of **X**, respectively. A complex 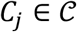 is defined to be a *balanced complex* (BC) for network *G*, if **A**^j:^**v** ≡ 0 for all 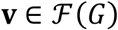. Given how incidence matrix **A** is constructed, this definition complies with the notion that the algebraic sum of fluxes entering and leaving *C_j_* must be always zero at any steady state.

### 2.3 Primal-dual formulations

To investigate whether a complex 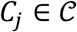 is a BC, one may form and solve the following two linear programs (LP)

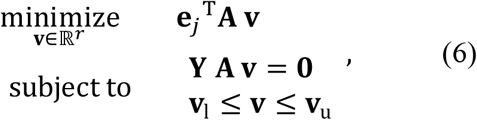

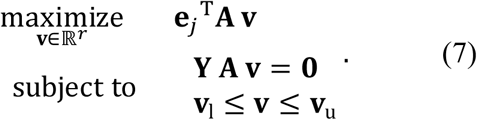

The definition implies that 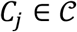 is a BC if and only if both above LPs have zero optimal values. The optimization problems (6) and (7) have a Lagrange dual of the following form [derivation in Section S1.2 of the Supplementary Information]

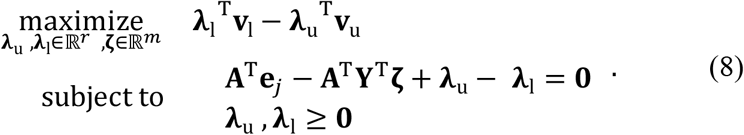

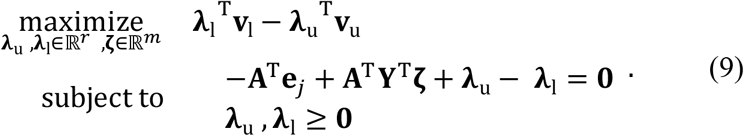

Clearly, given the linear constraints, strong duality holds and the duality gap is zero (Boyd & Vandenberghe, 2004); therefore, a complex 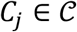 is a BC if and only if both Problems (8) and (9) have optimal values equal to zero.

## 3 Results and discussion

### 3.1 Generalization of trivial BCs

Taking into consideration the stoichiometry of a chemical reaction network, it is straightforward to see how some balanced complexes may emerge. A complex 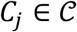 is *trivially-balanced* (a *trivial BC*), if there exist some species 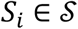 that appears only in *C_j_* and nowhere else in the network (Küken, et al., 2021). Given the structure of the stoichiometric map **Y**, it follows that **Y**^*i*:^ = e_j_ ^T^. Hence, the steady state equation **Y A v** = **O** immediately yields e_j_ ^T^**A v** ≡ 0,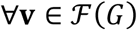.

One can readily generalize the idea behind the formation of a trivial BC to potentially arrive at a larger class of balanced complexes in the network. Intuitively, the vector **e**_*j*_ does not have to appear explicitly as a row in **Y**; all it takes is for ej to be a linear combination of rows in **Y**, that is

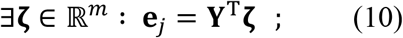

it then follows that **e**_*j*_ ^T^**A v** = **ζ**^T^**Y A v** = 0,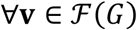. We remark that Eq. (10) can potentially explain a higher number of balanced complexes in the network, and it fully contains all the trivial BCs.

One may take another step and further generalize this notion to potentially identify an even larger class of BCs. Note that all BCs predicted by Eq. (10) have purely stoichiometric grounds, derived from stoichiometric map **Y**. However, the topological structure of a network, pinpointed by matrix **A**, can also play a role in the formation of BCs. Let us next consider a simple example, in which some species 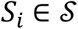 makes a unimolecular appearance in complexes 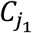 and 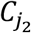, but is absent elsewhere in the network. Assume additionally that 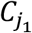, 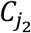 form a linkage class with another complex, namely 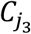. The situation is shown in Fig. 1.

**Fig. 1.**
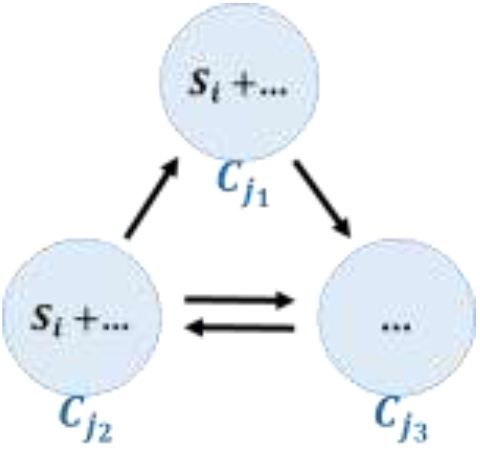
The topology contributing to the formation of a BC. A small linkage class of a network comprising three complexes is shown. The species S_*i*_ is assumed to be present in complexes 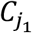 and 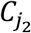, but nowhere else in the network.

On the one hand,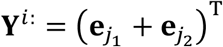 and the steady state condition for species *S*_*i*_ imply that 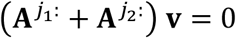,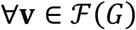 on the other hand, the closedness of the linkage class requires that 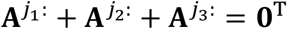. Thus, it follows that 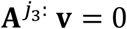, 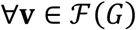, that is, 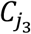 is a BC. Note that the balancing of 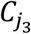 has nothing to do with the species in it, but with its connection to other complexes.

Formalizing this generalization relies on integrating the linkage structure of the network, which is closely interlinked with the left nullspace of the incidence matrix **A**, into Eq. (10). Let *G* be a closed network composed of *ℓ* linkage classes *L*^1^, …,  *L*_*ℓ*_. The left nullspace of **A**, i.e. ker **A**^T^ has a basis of the form {**u**_1_, …,  **u**_*l*_}, where 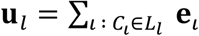, 1 ≤ *l* ≤ *ℓ*. Next, define the matrix **U** ∈ ℝ^n×*l*^ as **U** = [**u**_1_… **u**_*ℓ*_]. Now, let a complex ej have the following form

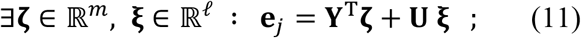

it follows that **e**_*j*_^T^**A v** = **ζ**^T^**Y A v** + **ξ**^T^**U A v** = 0, 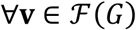. Eq. (11) shall be viewed as a generalization of Eq. (10), which can potentially account for a higher number of BCs in the network.

### 3.2 Factorizations of balanced complexes

Eq. (11), like Eq. (10), provides a sufficient condition: If there exist vectors ζ ∈ ℝ^*m*^, **ξ** ∈ ℝ^*ℓ*^ such that Eq. (11) holds for some **e**_*j*_, then complex 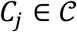 is a BC. This poses the following question: Does Eq. (11) account for all BCs in the network? We seek to determine whether all BCs are created as a result of an interplay of stoichiometry and the linkage structure or there exist other factors contributing to the formation of BCs. We are also interested in establishing the conditions under which Eq. (11) would become a necessary and sufficient condition for BCs.

The dual formulation in Eqs. (8) and (9) helps address these questions.

#### Theorem 1

A complex 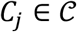 is a BC, if and only if there exist variables **ζ**_1_, **ζ**_2_ ∈ ℝ^*m*^, **λ**_u1_, **λ**_l1_, **λ**_u2_, **λ**_l2_ ∈ ℝ^*r*^, such that

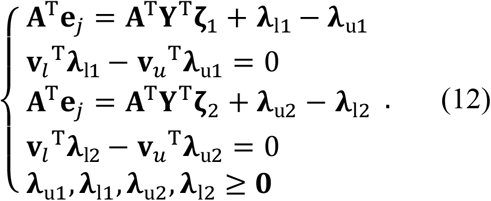

Eq. (12) implicitly relates the balanced complex not only to the stoichiometry but also to the upper- and lower bounds on reaction fluxes. It can actually be viewed as a generalization of Eq. (11), which brings into play the flux bound constraints as a third factor in the formation of BCs. The following result presents an intermediary formulation that elucidates the connection between Eqs. (11) and (12):

#### Theorem

Suppose the network is operating under a canonical flux regime. There exist variables **ζ**_1_, **ζ**_2_ ∈ ℝ^*m*^, ξ_1_, ξ_2_ ∈ ℝ^*l*^, **θ**_1_, **θ**_2_ ∈ ℝ^*n*^, such that

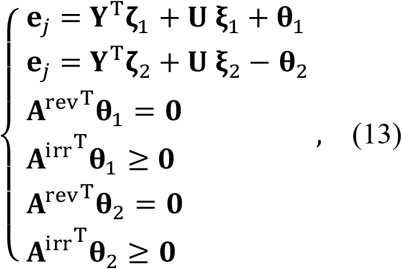

if and only if the complex 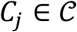 is a BC. By convention, **A**^irr^ denotes the columns of **A** corresponding to zero lower bounds in **v**_l_, and **A**^rev^ is the complementary block in **A**.

It is straightforward to see how Eq. (13) is a generalization of Eq. (11). In particular, any feasible solution of Eq. (11) for parameter values **ζ**, ξ corresponds to a solution of Eq. (13) with feasible parameter values **ζ**_1_ = **ζ**_2_ = **ζ**, ξ_1_ = ξ_2_ = ξ, **θ**_1_ = **θ**_2_ = **0**. Note that irreversibility patterns play a key role in distinguishing the solution sets of Eqs. (13) and (11). In addition, it is not difficult to show how Eq. (12) can be viewed as a generalization of Eq. (13). One only needs to define **λ**_l*t*_ = **A**^T^**θ**_*t*_, **λ**_u*t*_ = **O**; *t* = 1,2 [refer to Section S1.4 of the Supplementary Information for mathematical details].

### 3.3 Categorization of BCs

In Eqs. (10), (11), (13) and (12), we presented four different formulations that can explain the emergence of a BC potentially as a combined effect of distinct factors, namely stoichiometry, linkage structure, irreversibility patterns and flux bounds, respectively. Each “*factorization*” can be viewed as a feasibility problem; let us next define the *solution set* of each factorization as the set of all complexes 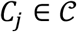, where vector **e**_*j*_ satisfies the feasibility problem for some parameter values. It is worth noting that the solution set of each factorization is a subset of the next one, in the abovementioned order of factorizations. Moreover, the solution set of Eq. (12) includes every existing BC in the network. This property paves the way to classify each BC of the network into one of the following four categories:

I. A complex 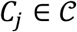 is called a *strictly stoichiometric* BC, if it has a factorization of the form Eq. (10), i.e. ∃**ζ** ∈ ℝ^*m*^: **e**_*j*_ = **Y**^*T*^**ζ**. This relation is referred to as a *strictly stoichiometric factorization* for the BC represented by vector **e**_*j*_.
II. A complex 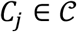 is called a *stoichiometric* BC, if it has a factorization of the form Eq. (11), i.e. ∃**ζ** ∈ ℝ^*m*^, **ξ** ∈ ℝ^*ℓ*^: **e**_*j*_ = **Y**^T^**ζ** + **U ξ**. This relation is referred to as a *stoichiometric factorization* for the BC represented by vector **e**_*j*_.
III. A complex 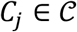 is called a *type-I nonstoichiometric* BC, if it has a factorization of the form Eq. (13), but no stoichiometric factorization. Eq. (13) is called a *type-I nonstoichiometric factorization* for the BC represented by vector **e**_*j*_.
IV. A complex 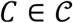 is called a *type-II nonstoichiometric* BC, if it has a factorization of the form Eq. (12), but no factorization of the form Eq. (13). Eq. (12) is called a *type-II nonstoichiometric factorization* for the BC represented by vector **e**_*j*_.

It follows directly from these definitions that any strictly stoichiometric BC/factorization is also a stoichiometric BC/factorization. Moreover, a balanced complex is labeled with the general term *nonstoichiometric BC*, if it has no stoichiometric factorization, regardless of whether it has an explicit factorization of the form Eq. (13) or an implicit factorization of the form Eq. (12).

### 3.4 Characteristics of distinct BC categories

Recent studies show that balanced complexes can play an important role in the reduction of metabolic networks and simplification of steady state analysis (Feliu & Wiuf, 2013; Küken, et al., 2021). The categorization of BCs into distinct classes not only helps identify the origin of the balancing in each case, but also enables one to study their properties. On the one hand, this equips us with an insight into how certain modification of a network may affect existing BCs. On the other hand, it may demonstrate if and how modeling inaccuracies may lead to a detection failure for certain existing BCs.

The strictly stoichiometric BCs constitute the most trivial class, whose formation relies merely on the network stoichiometry. This simplicity gives rise to an intrinsic form of robustness: As long as network complexes are unaltered, the balancing property of strictly stoichiometric BCs remains preserved, regardless of any potential changes in the network, e.g. even if the reactions in these networks are altered (referred to as different enzymatic regimes). In this case, the formation of strictly stoichiometric BCs roots solely in the steady state property for the species.

For the more general class of stoichiometric BCs, their formation not only relies on the network stoichiometry, but also partly on a topological feature, namely the linkage structure. They are affected neither by the irreversibility patterns in the network nor by the constraints on reaction rates. However, they may be altered under different enzymatic/catalytic regimes. Their balancing property is rooted not only in the steady state property but also in the conversion patterns (i.e. reaction structure) of the network.

The formation of nonstoichiometric BCs is a more sophisticated and at the same time more interesting phenomenon. They often emerge as a combined effect of several network features, including the stoichiometry and the linkage structure, but also irreversibility patterns as well as upper- and lower bounds on fluxes. In other words, these BCs carry information in the form of an equality constraint on reaction fluxes, which is not encoded in the steady state equation **Y A v** = **0**.

In a canonical flux regime, i.e. when irreversible reactions have a zero lower bound on reaction rates, all nonstoichiometric BCs are in the type-I category. As their factorization suggests, their emergence relies specifically on the irreversibility patterns and not on the exact values of the flux bounds. This makes the detection of these BCs insensitive to potential modeling errors in the form of inaccuracies in flux bounds, even when they appear in a non-canonical flux regime. However, BCs in this category have the following undesirable property:

#### Proposition 3

Let a network *G* contain a type-I nonstoichiometric BC. There exist at least two blocked irreversible reactions in the network.

It follows that once all blocked reactions are removed, no type-I nonstoichiometric BCs may exist in the network. In fact, the removal of such reactions changes the linkage structure of the network such that all type-I nonstoichiometric BCs in the original network appear as stoichiometric BCs in the reduced network.

Type-II nonstoichiometric BCs do not automatically require a number of network reactions to be blocked in the steady state flux set. By contrast, they require a number of reactions to be fixated at a nonzero upper-or lower bound, as the following statement demonstrates.

#### Proposition 4

Let *G* be a network all blocked reactions of which have been removed. Suppose *G* contains a nonstoichiometric BC. There exist at least three reactions in *G*, which are fixated at a corresponding nonzero lower-or upper bound, for all 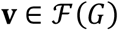.

More specifically, it can be shown that the existence of a type-II nonstoichiometric BC implies at least one irreversible reaction is fixated at a positive lower bound. It is not surprising that such BCs only emerge in non-canonical flux regimes. However, an undesirable feature with type-II nonstoichiometric BCs is that their factorization relies explicitly on the values of upper-and lower bounds, which may make their detection sensitive to modeling assumptions.

### 3.5 Toy examples

Distinct classes of balanced complexes identified in this study are practically observed in all sorts of small and large networks. However, here we present small networks specifically designed for illustrative purposes. The fact that the chemistry in these toy networks may seem unrealistic or may not reflect real-world mechanisms should not be viewed critically; they were constructed with the idea of providing simple illustrations involving very few species.

With that in mind, let us next consider the toy network on Fig. 2.

**Fig. 2.**
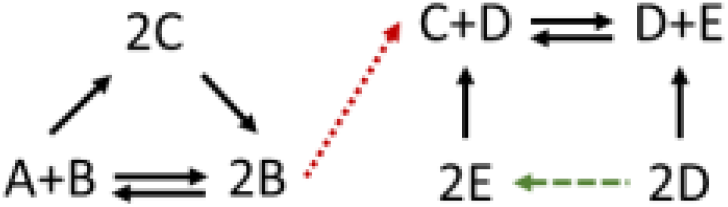
Stoichiometric BCs in a toy network. The conversion diagram depicts a base network with *m* = 5 species, *n* = 7 complexes, and *r* = 6 base reactions, two of which are reversible. The network has four BCs, which will remain balanced even if one adds the dashed reaction (highlighted in green). However, once one adds the dotted reaction (highlighted in red), only two of these complexes remain balanced.

Let us first only consider the base network in Fig. 2, which excludes the dotted and dashed reactions. It is trivial to show that (*A* + *B*) and (2*B*) have strictly stoichiometric factorizations. The two complexes (2*C*) and (*C* + *D*) are stoichiometric BCs but not strictly stoichiometric ones. All four complexes will remain balanced, if one changes the irreversibility of reactions arbitrarily or if the dashed reaction is added to the network. However, as soon as one adds the dotted reaction to the network, the linkage structure will be altered. As a result, the strictly stoichiometric BCs (*A* + *B*) and (2*B*) will still be balanced, but the stoichiometric BCs (2*C*) and (*C* + *D*) will not remain balanced.

Nonstoichiometric BCs are more complex phenomena that are expected to emerge more frequently in larger networks, and, thus, are more challenging to illustrate in toy networks. To demonstrate an example, let us consider the conversion diagram in Fig. 3.

**Fig. 3.**
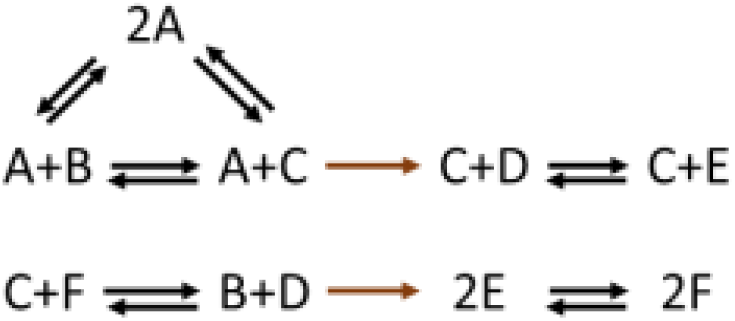
A nonstoichiometric BC in a toy network. The conversion diagram depicts a network with *m* = 6 species, *n* = 9 complexes, and *r* = 8 reactions. The network has one balanced complex (2*A*) which is a type-I nonstoichiometric BC. As predicted by Proposition 3, the two irreversible reactions in this network (highlighted in brown) are both blocked at steady state.

The network in Fig. 3 contains no stoichiometric BCs. Let us sort the species by alphabetical order and sort the complexes as follows: (*C*_1_, …,  *C*_9_) = (*A* + *B*, *A* + *C*, *C* + *D*, *C* + *E*, *C* + *F*, *B* + *D*, 2*E*, 2*F*, 2*A*). Given the linkage structure of this network, let us define the basis matrix **U** as follows

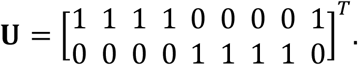

It can be shown that the complex indexed by vector **e**_9_ has a type-I nonstoichiometric factorization with the following parameters

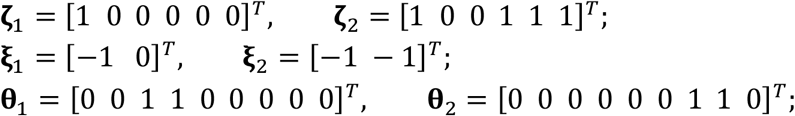

It follows that *c*_9_ is a type-I nonstoichiometric BC for the network. It can also be shown that the two irreversible reactions (shown in brown) are blocked at steady state, which is consistent with the prediction of Proposition 3. Note that removing the two blocked reactions would alter the linkage structure of the network, as a result of which the complex *C*_9_ = 2A would become a stoichiometric BC for the reduced network.

It is worth noting that changing the irreversibility patterns elsewhere in the network does not affect the balancing property; however, *C*_9_ would not be balanced anymore if one modifies either of the two reactions in brown. It is precisely the irreversibility of these two reactions in this particular constellation that makes *C*_9_ a balanced complex.

### 3.6 Presence of the different categories of BCs in metabolic models

To show the prevalence of the proposed categorization of BCs in real-world networks, we categorize balanced complexes identified in twelve genome-scale metabolic networks of organisms from all kingdoms of life (Küken, et al., 2021). Küken et al. removed blocked reactions from the networks prior to the identification of BCs, which results in the absence of type-I nonstoichiometric BCs. Our results indicate that all remaining categories of BCs, namely: strictly stoichiometric, stoichiometric, and type-II nonstoichiometric, are found across all analyzed genome-scale networks. We find that stoichiometric BCs form the largest group of BCs with 59% to 94% of all BCs identified, except for models of *A. thaliana*, *M. musculus*, *N. pharaonis* and *P. putida* where stoichiometric BCs could not be found or comprise below 1.5% of all BCs (Fig. 4). In contrast, the class of strictly stoichiometric BCs is present in all twelve analyzed models, with 6% to 94% (on average 26%) of BCs falling into this category. Type-II nonstoichiometric BCs could be identified in the metabolic models of *A. thaliana* (6%), *M. musculus* (66%), *N. pharaonis* (93%) and *P. putida* (90%). Interestingly, models including type-II nonstoichiometric BCs, were found to include less than 1.5% stoichiometric BCs. Our analysis of metabolic networks from organisms of kingdoms across life shows the prevalence of all proposed factors (stoichiometry, linkage structure and flux bounds) for the formation of BCs in biological networks.

**Fig. 4.**
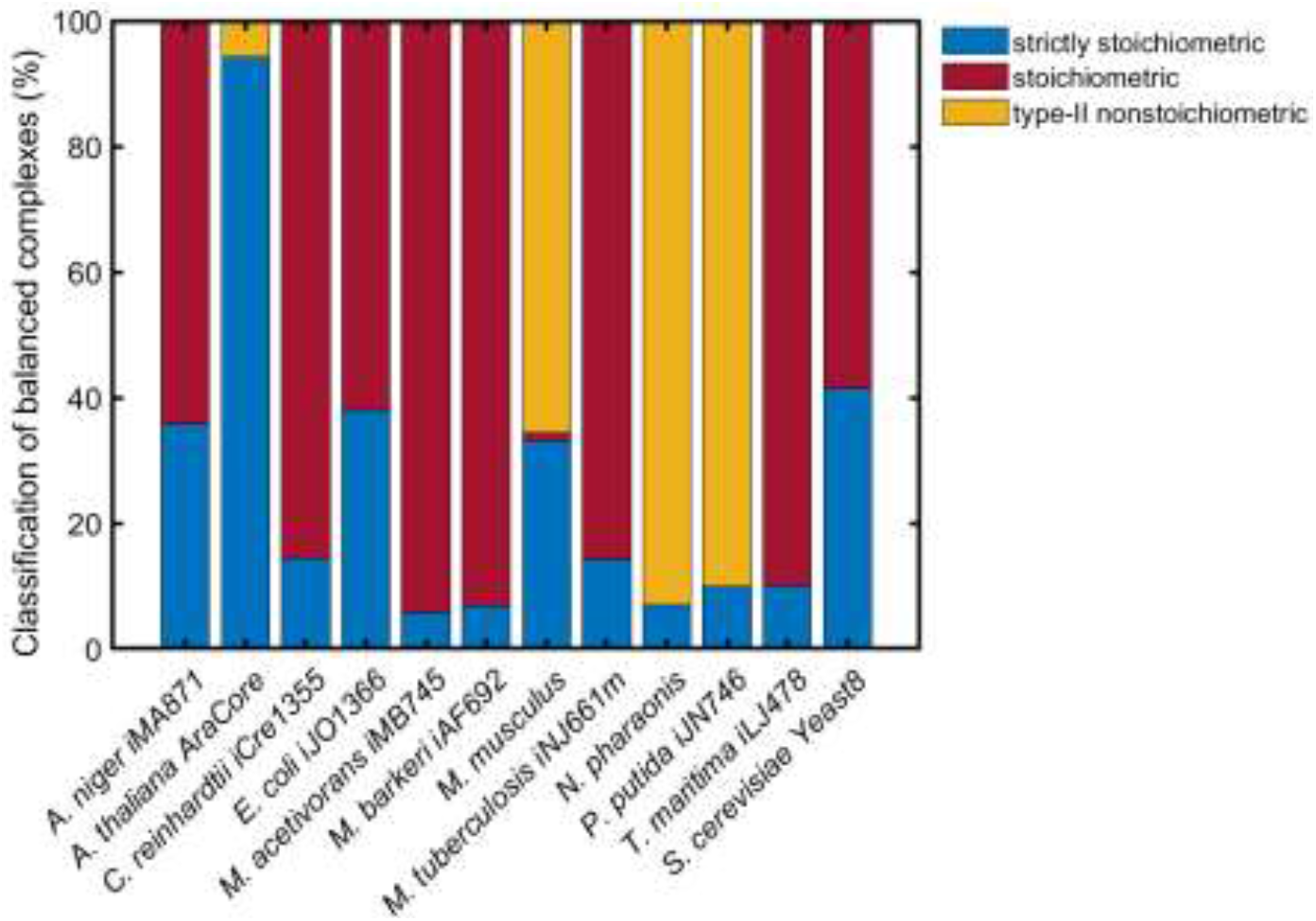
Classification of balanced complexes into strictly stoichiometric, stoichiometric and type-II nonstoichiometric. Balanced complexes from twelve genome-scale models of organisms across kingdoms of life are classified and the percentage of model BCs that fall into the respective class is presented. Type-I nonstoichiometric BCs are not present in the analyzed models since blocked reactions were removed from the networks prior to analyses. For model references, see Supplementary Information.

## 4 Concluding remarks

The nonlinearity and dimensionality of steady state equations makes it extremely difficult to study the properties of metabolic networks. While closed-form solutions seem evasive in most cases, recent approaches exploit balanced complexes as a means to reduce metabolic networks and facilitate steady state analysis. However, the underlying mechanisms leading to the formation of balanced complexes have not been studied so far. This work investigates this question and identifies stoichiometry, linkage structure, irreversibility patterns and flux bounds as the main driving factors behind this phenomenon.

The analysis enables us to classify balanced complexes into distinct stoichiometric and nonstoichiometric categories based on their origins and to study their properties. Formulation of the problem as a linear feasibility program provides a computationally efficient tool that is applicable to even large-scale metabolic networks. It is a fascinating aspect of these results that they are not obtained under any specific kinetic assumptions, which suggest they enjoy a ubiquity and are unperturbed by kinetic approximations. Indeed, examination of several well-established metabolic models demonstrates that all these categories are present in distinct metabolic networks across diverse kingdoms of life.

## Supplementary Information

The full derivation of the dual problems and all mathematical formulations, as well as detailed proofs for the theorems and propositions in the text are provided in <u>Supplementary Information [link]</u>. In addition, we provide information on the analyzed genome-scale metabolic networks and computational procedure of BC classification.

## 5 S1 Mathematical Analysis of Balanced Complexes

### 5.1 S1.1 Definitions and motivation

Let 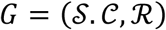 be a chemical reaction network with stoichiometric map **Y** and complex-to-reaction incidence matrix **A**. The steady state flux set for *G* is defined as follows

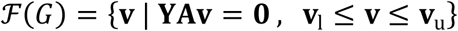

where **v**_l_ and **v**_u_ denote lower-and upper bounds for flux through reactions, respectively.

A complex 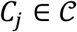 is referred to as a balanced complex (BC), if for all flux vectors in 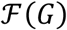 the total flux entering this complex is equal to the total flux leaving this complex. This is equivalent to

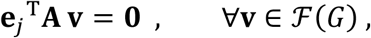

where **e**_j_ is the vector with a unit value for the *j*^th^ entry and zero values elsewhere.

For any 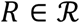, we say *R* is a blocked reaction in *G*, if for all the feasible set distributions 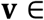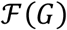, there is zero flux through *R*, that is, 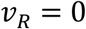. This is a commonly occurring phenomenon in metabolic networks, especially in scenarios when flux bounds and/or optimization of particular objectives are imposed. Similarly, we say a reaction 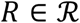 is fixated at some flux value *f*, if for all flux distributions 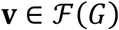, the flux through *R* is unchanged, namely, 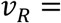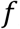, 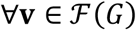. Clearly, any blocked reaction is one fixated at zero.

Looking at the stoichiometry of a chemical reaction network, it is easy to see how balanced complexes may emerge. In the most trivial case, any complex comprising a species that appears nowhere else in the network must be balanced at any steady state, thus a BC. Note that this case amount to finding a single row of the stoichiometric map **Y** –as a binary indexing vector for a complex- that defines a balanced complex.

One can readily generalize this idea to come up with more balanced complexes for the network. Suppose a vector **e**_*j*_ can be expressed as a linear combination of rows of the stoichiometric map **Y**, that is

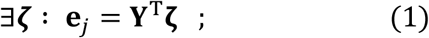

it immediately follows that

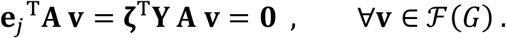

One can further expand the idea by taking into account the left nullspace of the incidence matrix **A**. Let *G* be a closed system; the left nullspace of **A** has a simplified structure: If the CRN has a connected graph, then the nullspace is the one-dimensional subspace defined as span of basis vector, 〈**1**〉. This yields

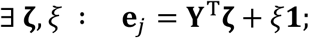

If the CRN is not connected, then the nullspace has basis vectors of the same structure, but for separate linkage classes. Let us assume *G* consists of *ℓ* linkage classes, denoted by *L*_1_, *L*_2_, …,  *L*_*ℓ*_. With each linkage class *l*, for *l* = 1, …,  *ℓ*, one can associate a vector 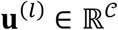 defined as follows

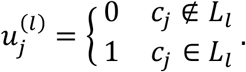

The columns of the matrix 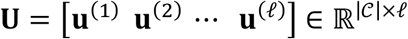 form a basis for the left nullspace of **A**. Now suppose a vector **e**_*j*_ can be expressed as follows

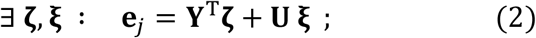

it is easy to see that

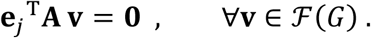

Therefore, **e**_*j*_ represents a balancing relation for *G*.

We refer to Eq. (2) as a stoichiometric factorization for the indexing vector **e**_*j*_. Any BC that can be parameterized in the form of Eq. (2) is referred to as a stoichiometric BC.

Here come the key questions we want to address next: Are these the only forms of balanced complexes that may emerge in a network? In other words, are all balanced complexes in a network simply a result of the stoichiometric-and linkage structure? Could other constraints on a network, such as lower and upper bounds on flux levels, also play a role in creating some other balanced complexes?

### 5.2 S1.2 The primal-dual formulation

To answer these questions, we first formulate the balancing problem as a pair of optimization problems as follows

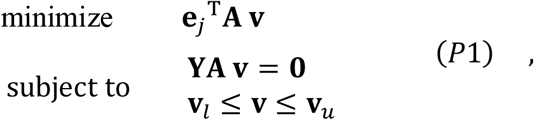

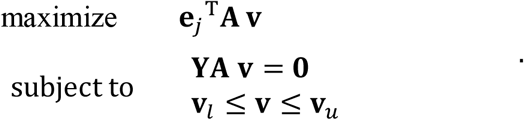

For the problem formulation to conform to standard conventions in convex optimization, we simply replace the 2^nd^ optimization problem by an equivalent minimization problem as follows.

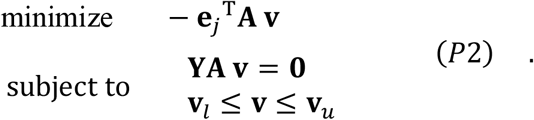

A vector **e**_*j*_ represents a balanced complex, if and only if both optimization problems *P*1 & *P*2 have an optimal value of zero.

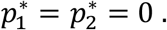

Any BC has to satisfy this criterion.

To further analyze which vectors **e**_*j*_ may satisfy this criterion, we will transform these optimization problems into Lagrange dual forms. It is well-established that strong duality will hold for these convex problems with linear constraints [1], hence the optimal duality gap is zero. Therefore, instead of directly solving *P*1 and *P*2, one can try to solve the dual problems.

To derive the dual problem for *P*1, let us first define the Lagrangian as follows

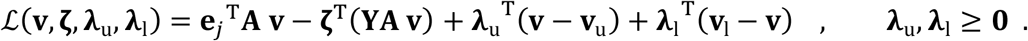

The Lagrange dual function is then defined as follows

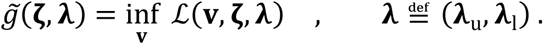

Since the Lagrangian is an affine function of **v**, it is unbounded from below, hence its infimum value is in general −∞, unless the slope of the affine function is zero, that is

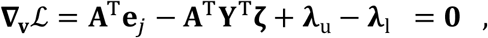

in which case, the Lagrange dual function would be simplified to the following function

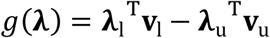

Hence, the dual problem is

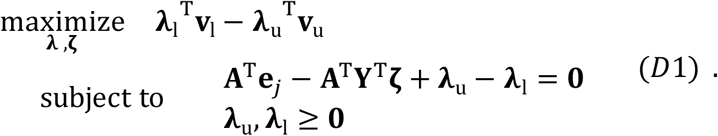

Similarly, one can process the optimization problem *P*2 to arrive at the following dual formulation

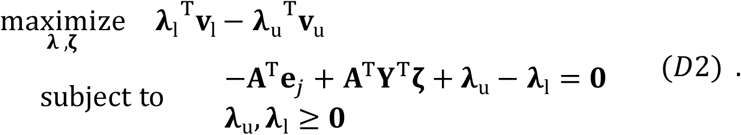

For **e**_*j*_ to represent a BC, both problems D1 and D2 must have zero optimal values, i.e.

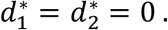

It is worth noting that, while the primal problems *P*1 and *P*2 have a slightly different objective but share the same constraints, the dual problems share the same objective function while having a slightly different feasible set.

### 5.3 S1.3 Analyzing the dual problems(s) in a canonical flux regime

#### 5.3.1 S1.3.1 Analyzing the dual objective

The objective function for the dual problems can be written as

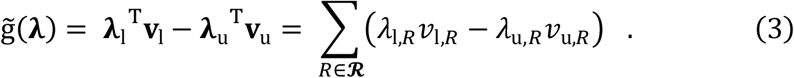

To simplify this objective function, let us assume the CRN operates in a *canonical flux regime*, that is the bounds on reaction fluxes are standardized as follows

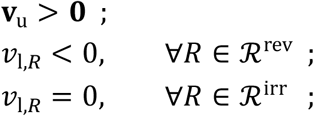

where 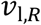 denotes the *R*^th^ entry of vector **v**_l_, corresponding to reaction 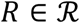 In a canonical regime, the irreversible reactions are exactly those corresponding to zero entries of **v**_l_, while the rest of entries in **v**_l_ must be negative and correspond to reversible reactions.

In a canonical regime, it is easy to verify that all terms on the right hand side of Eq. (3) are nonpositive, given the dual feasibility conditions **λ**_u_, **λ**_l_ ≥ **0**. For the sum of all those terms to equal zero, every single term in the RHS must be zero. At any optimal point, we must have

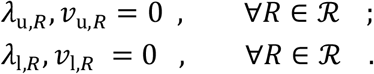

This, in turn, yields,

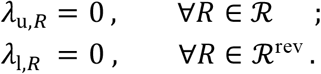

Therefore, for the complex balancing to hold, out of all entries in **λ**_u_, **λ**_l_, only those associated with lower bounds of irreversible reactions can take arbitrary values (though they must be nonnegative due to dual feasibility). The rest of entries in **λ**_u_, **λ**_l_ must all be zero.

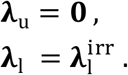

Note that **λ**_u_, **λ**_l_ having the above form is a necessary condition for the dual problem(s) to admit feasible solution(s) with an objective value of zero. Moreover, if such a solution is found, then it is clearly an optimal point.

It is also worth noting that under a canonical flux regime, the upper bounds on flux levels (as well as negative lower bounds on flux through reversible reactions) have no impact on the formation of balanced complexes. Hence, as far as the emergence of balanced complexes is concerned, it makes sense to just ignore those upper bounds on fluxes and simply study the resulting steady state flux cone.

Moreover, for some **e**_*j*_ to represent a BC, the dual variables must also satisfy the other constraints in dual problems *D*1 and *D*2.

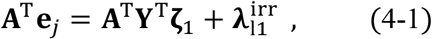

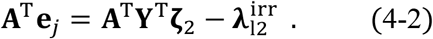

Both the above equations must hold, for the dual problems *D*1 and *D*2 to both have zero optimal values, that is, 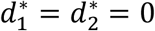.

Therefore, Any vector **e**_*j*_ which satisfies the above two equations for arbitrary parameter values **ζ**_1_, **ζ**_2_ and 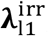,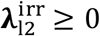 provides a zero optimal solution for the primal optimization problem and hence, represents a balanced complex for the given CRN. In other words, this can be seen as a necessary and sufficient condition, under a canonical flux regime.

#### 5.3.2 S1.3.2 Relation to stoichiometric factorization

Let us further study the two equalities in Eq. (4). First of all, both vectors 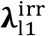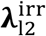 must lie in the range of **A**^T^, that is, the row space of **A**. Hence, we can introduce a change of variables

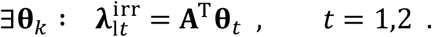

Given the discussion in Section S1.3.1, each new variables **θ**_*t*_ must satisfy the following constraints,

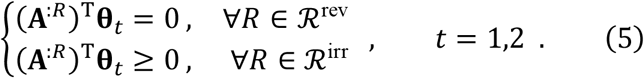

Therefore, one can replace the parameters 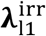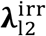 in Eq. (4) with their equivalents in terms of **θ**_1_ and **θ**_2_, while integrating extra constraints [of dual feasibility] given in Eq. (5).

Using the new parameter **θ**_1_, Eq. (4-1) can be written as

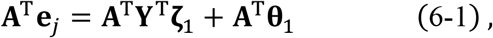

in which, **ζ**_1_ and **θ**_1_ are independent parameters that are only constrained to satisfy Eq. (5). The general solution for **e**_*j*_ in Eq. (6-1) will be of the form

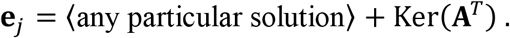

Assuming the CRN is closed, Ker(**A**^*T*^) has the well-known structure discussed in Section S1.1. Therefore, from Eq. (4), one obtains the following general expressions for **e**_*j*_

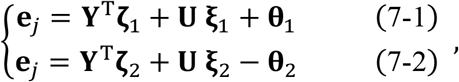

for independent parameters **ζ**_1_, **ζ**_2_, **ξ**_1_, **ξ**_2_, **θ**_1_, **θ**_2_, subject to the constraints expressed in Eq. (5).

A special case for **e**_*j*_ may be obtained by setting **θ**_1_ = **θ**_2_ = 0. Note that this choice of parameter values also satisfies the constraints in Eq. (5). With **θ**_1_ = **θ**_2_ = 0, Eq. (7) reduces to two identical equations of the form

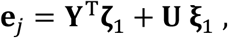

which is exactly the stoichiometric factorization presented in Eq. (2). Therefore, the factorization in Eq. (7), obtained by solving the dual optimization problems, can be viewed as a generalization of the stoichiometric factorization.

In a canonical flux regime, a vector **e**_*j*_ is said to represent a nonstoichiometric BC, if it has a factorization corresponding to Eq. (7), but no stoichiometric factorization [corresponding to Eq. (2)]. For reasons that will become clear soon, any such a BC is referred to as a *type-I nonstoichiometric BC*.

Now, if one partitions the incidence matrix **A** into reversible and irreversible blocks as in **A** = [**A**^rev^ **A**^irr^], then the constraints in Eq. (5) can be rewritten in the following simpler block form

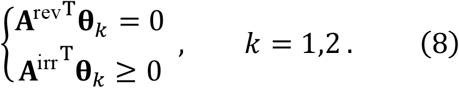

#### 5.3.3 S1.3.3 The factorization of BCs in a canonical flux regime

Assuming the system operates in a canonical flux regime, the discussion above can serve as a proof for the following statement.

##### Theorem S1

Suppose the network is operating in a canonical flux regime. There exist variables 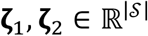, **ξ**_1_, **ξ**_2_ ∈ ℝ^*ℓ*^,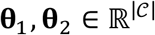, such that

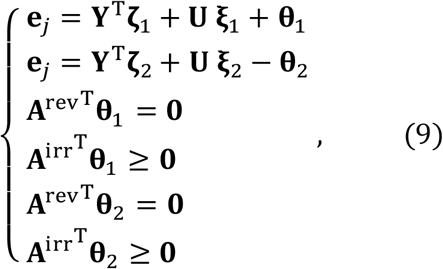

if and only if the complex 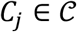 is a BC.

Due to the linearity of the above factorizations and constraints, Eq. (9) can serve as a linear feasibility problem, with variables **ζ**_1_, **ζ**_2_, **ξ**_1_, **ξ**_2_, **θ**_1_, **θ**_2_, to check whether any given complex 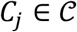 is a BC of the network. In view of the highly efficient computational tools available for LP problems, this provides an alternative approach for finding balanced complexes. What comes on top of it as a bonus is the fact that it allows one to systematically identify the underlying factors contributing to the formation of any BC.

Let us further reflect on these constraints. Out of all variables in the factorization, only **θ**_1_, **θ**_2_ are constrained by the following constraints, and **ζ**_1_, **ζ**_2_, **ξ**_1_, **ξ**_2_ are not. Hence, one may transform the constraints in Eq. (9) into the following-more compact-form

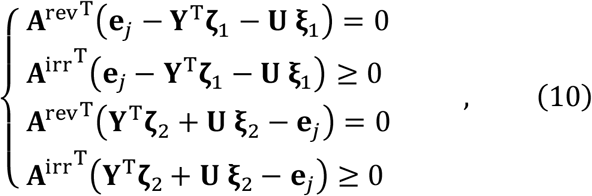

which can replace Eq. (9) in the LP framework. This may be more practical in terms of implementation, but not necessarily in terms of clarification, because it fails to provide an explicit factorization of **e**_*j*_, unlike Eq. (9).

##### 5.3.4 S1.3.4 Formation of type-I nonstoichiometric BCs

The following statement sheds light on the properties on type-I nonstoichiometric BCs and how they are formed.

###### Proposition S2

<u>Proof</u>: Let **e**_*j*_ represent the type-I nonstoichiometric BC. It immediately follows from the definition that we must have

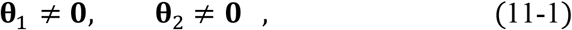

as well as,

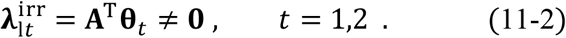

The balancing relation itself implies that

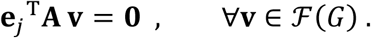

Replacing **e**_*j*_ by its factorization equations yields

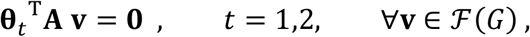

thsi is,

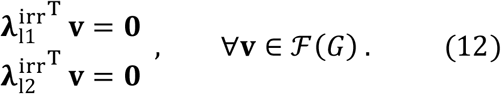

Note that

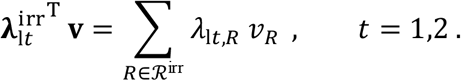

Having the abovementioned structure of the dual variables 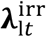, *t* = 1,2, and knowing that 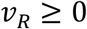,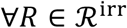 it follows from Eq. (12) that

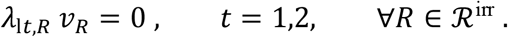

Taking into consideration that Eq. (11-2) implies 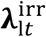 has at least one positive entry for both *t* = 1,2, it follows that for **e**_*j*_ to represent a nonstoichiometric BC, at least one irreversible reaction must be blocked.

Moreover, the two vectors 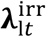, *t* = 1,2 cannot be collinear; otherwise one can cancel out those terms in Eq. (4), which yields a stoichiometric factorization for the BC, which would be a contradiction. Therefore, there exist at least two blocked reactions in the network.∎

**Corollary S3**. Let *G* be a network operating under a canonical flux regime, all blocked reactions of which have been removed. Then all balanced complexes of *G* have stoichiometric factorizations.

#### 5.4 S1.4 Analyzing the dual problem(s) for general flux bounds

##### 5.4.1 S1.4.1 Derivation of an equivalent LP

Next, we consider the network under general -not necessarily canonical- flux bounds, in which some irreversible reactions may also take positive lower bounds on flux levels. Irrespective of the flux bounds, for a vector **e**_*j*_ to represent a BC, the dual optimization problems *D*1 and *D*2 must take zero optimal values.

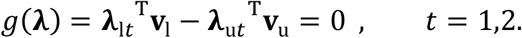

Here is what contrast this case from the canonical flux regime: Some terms in the dual objective may now take positive values; hence we are not in a position to further simplify the objective function by asserting that every term on the RHS has to be zero, as we did in Section S1.3.1.

A complex represented by vector **e**_*j*_ is balanced if and only if both dual problems D1 and D2 have zero optimal values. Now, we represent the equivalent of Theorem S1 for the more general case:

###### Theorem S4

A complex 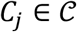 is a BC, if and only if there exist variables 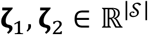, 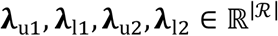 such that

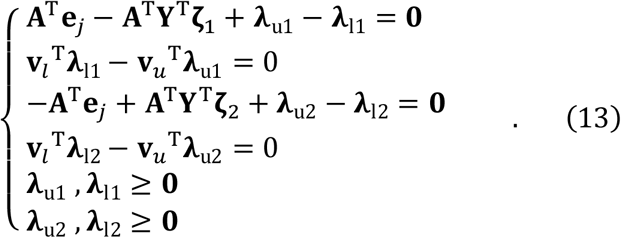

<u>Proof</u>: let us momentarily take a break from directly addressing optimality conditions for D1 and D2, and instead try to seek feasible variables that set the corresponding dual objectives to zero. Hence, we now seek variables 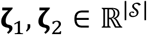,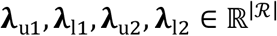 that satisfy Eq. (13).

Note that (**ζ**_1_, **λ**u_1_, **λ**_l1_) gives a feasible point for *D*1, while (**ζ**_2_, **λ**_u2_, **λ**_l2_) gives a feasible point for *D*2. Therefore, if one denotes the optimal (maximal) values of *D*1 and *D*2 by 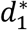, 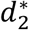 respectively, the feasibility of these points implies that

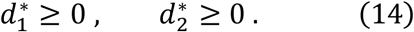

On the other hand, 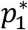 is the minimum of some function (**e**_*j*_ ^T^**A v**) on the steady state flux set, while 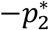 is the maximum of the same function on the same set. Hence, 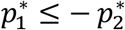. Therefore

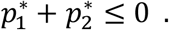

As a result, the general properties of the duality [1] ensure that

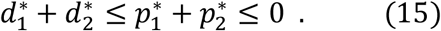

Taking into account Eqs. (13) and (15), it follows that

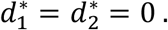

Therefore, even though we did not explicitly enforce optimality in Eq. (13), by the virtue of Eq. (5), any set of variables that satisfies Eq. (13) automatically provides optimality, hence it translates to solutions for the dual optimization problems *D*1 and *D*2, with an achieved optimal value of zero.

On the other hand, if vector **e**_*j*_ represents a BC, it has to lead to zero optimal values for the dual problem(s), hence there exist variables 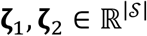, 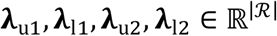that satisfy Eq. (13). ∎

As a result, in the general –not necessarily canonical- case, Eq. (13) replaces the duality problems, and may serve as a linear feasibility problem to check whether a complex **e**_*j*_ is a BC.

##### 5.4.2 S1.4.2 Nonstoichiometric balancing with general boundary constraints

Given the two equalities containing **e**_*j*_ in Eq. (13), it follows that the terms **λ**u_1_ − **λ**_l1_ and **λ**_*u*2_ − **λ**_l2_ must lie in the rowspan of **A**; therefore, similar to Eq. (7), one can introduce new variables **θ**_1_ and **θ**_2_, as follows

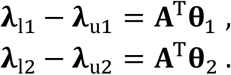

Using new variables **θ**_1_ and **θ**_2_, one would arrive at the same factorization structure as in Eq. (7) for the balanced complex **e**_*j*_, which could pave the way for labeling of **e**_*j*_ as either a stoichiometric or nonstoichiometric balancing relation. However, note that the other constraints in Eq. (9) do not hold generally, under non-canonical flux regimes. In fact, the solution set of Eq. (9) is a subset of the solution set of Eq. (13), for which those additional constraints also hold.

Instead of using new variables **θ**_1_ and **θ**_2_ to identify nonstoichiometric BC, one can resort to the following equivalent definition: A vector **e**_*j*_ represents a *nonstoichiometric BC*, if it has an implicit factorization corresponding to Eq. (13), but no factorization of the form Eq. (2).

As we have seen with Proposition S2 and Corollary S3, blocked irreversible reactions play a key role in the formation of (type-I) nonstoichiometric BCs in a canonical flux regime. It follows that they can play exactly the same role in networks operating under general flux bounds. However, one might ask the question: Can other factors also lead to the formation of BCs under general –not necessarily canonical- flux bounds?

To answer this question, let us suppose all blocked reactions have been removed from the network. It follows from Corollary S3 that no type-I nonstoichiometric BC may exist in this network. We aim to investigate whether possibly different forms of nonstoichiometric BCs may emerge in this network. Let us define 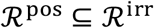 as the set of irreversible reactions with positive lower bounds on flux levels. Let us now assume there exists a nonstoichiometric BC in this network, represented by vector **e**_*j*_. From the definition of a nonstoichiometric BC, it follows that

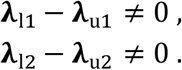

Moreover, it can be shown that the above two vectors cannot be collinear. Otherwise the equality

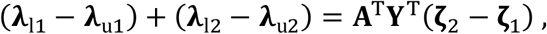

would mean that both (**λ**_l*t*_ − **λ**_u*t*_), *t* = 1,2 can be replaced by a scalar factor of **A**^T^**Y**^T^(**ζ**_2_ − **ζ**_1_), hence **e**_*j*_ would have a stoichiometric factorization, which is a contradiction. Therefore, the two vectors (**λ**_l*t*_ − **λ**_u*t*_), *t* = 1,2 are linearly independent. The other constraints in Eq. (13),

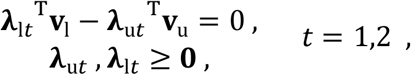

alongside **v**_u_ > **0** imply that **λ**_u*t*_^T^**v**_u_ ≥ 0, **λ**_l*t*_^T^**v**_l_ ≥ 0, *t* = 1,2; and we must also have

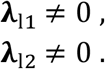

In addition, it is easy to show [by contradiction]

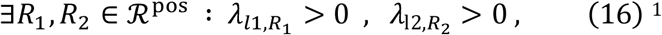

in which, 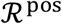 denotes the set of (irreversible) reactions with a positive flux lower bound.

From the definition of the balanced complex, we know that **e**_*j*_^T^**A v** = **0**,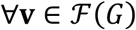.

Replacing **e**_*j*_^T^**A** = (**A**^T^**e**_*j*_)^T^ by its equivalent from Eq. (13) then yields

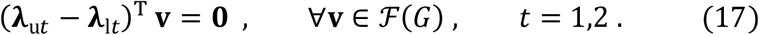

Therefore, existence of a nonstoichiometric balancing relation in the network implies the existence of at least two independent linear couplings of reaction fluxes.

In addition, via combining Eq. (17) with Eqs. (13-2) and (13-4), one obtains

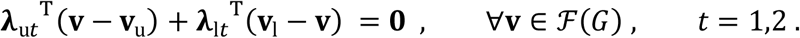

However, we know that

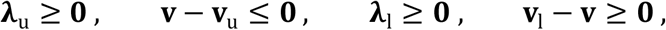

Therefore, all reactions for which at least one corresponding entry of **λ**_u1_, **λ**_u2_, **λ**_*l*1_, **λ**_*l*2_ is nonzero, must be fixated at their respective lower/upper bounds for all values of **v** in the steady state flux set. In particular, Eq. (16) ensures that for at least one irreversible reaction with a positive lower boundis fixated at the minimum for all 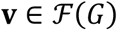

###### Proposition S5

Let *G* be a network all blocked reactions of which have been removed. Suppose *G* contains a nonstoichiometric BC. There exist at least one irreversible reaction with a positive lower bound 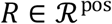, such that 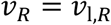,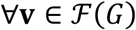.

The following statement follows immediately from Proposition S5.

###### Proposition S6

Let *G* be a network all blocked reactions of which have been removed. Suppose *G* contains a nonstoichiometric BC. There exist at least three reactions in *G*, which are fixated at a corresponding [nonzero] lower-or upper bound, for all 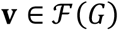.

<u>Proof</u>: From Eq. (16), we know that 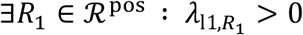 and the reaction *R*_1_ is fixated at the corresponding lower bound. From the BC constraints in Eq. (17), we have 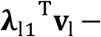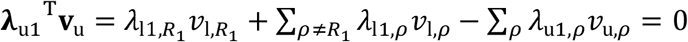. Since 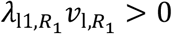 it follows that

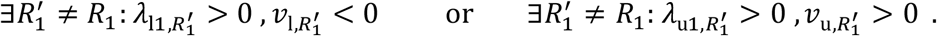

Therefore, there exist at least two distinct reactions 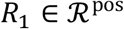,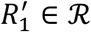 fixated at their corresponding flux bounds.

With the same logic, the other constraint 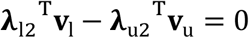 implies that there exist at least two distinct reactions 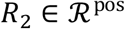,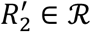fixated at their corresponding flux bounds. Now, if there exist any other reaction 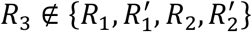 for which the strict inequality 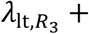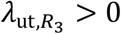 holds for some *t* ∈ {1,2}, then we have found the three fixated reactions and the proof is complete.

If that is not the case, then we cannot have both *R*_1_ = *R*_2_ and 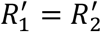; otherwise the two vectors (**λ**_l1_ − **λ**_u1_), (**λ**_l2_ − **λ**_u2_) would become collinear, which is a contradiction. Therefore, there exist at least three reactions fixated at nonzero upper- or lower bounds.∎

##### 5.4.3 S1.4.3 Type-I vs. type-II nonstoichiometric BCs

Let us remind that in networks operating under a canonical flux regime, any BC encoded by vector **e**_*j*_ must satisfy the factorization scheme presented in Eq. (7). We showed that for a nonstoichiometric BC to exist, at least two irreversible reactions must be blocked at steady state. It can also be shown that those blocked reactions cannot lie within any strong linkage class of the network, but each must be a “bridge” reaction that connects distinct strong linkage classes within a linkage class 2. We refer to this class of nonstoichiometric BCs as <u>*type-I*</u>.

It is not surprising that type-I nonstoichiometric BCs can also emerge in non-canonical flux regimes. To seek type-I nonstoichiometric BCs under general flux bounds, one can basically apply the same formulation as Eq. (9). There is, however, a minor caveat due to the potential existence of positive lower bounds on fluxes. Eq. (9) was introduced in the context of canonical flux bounds, where **A**^irr^ denoted exactly those columns of **A** corresponding to zero lower bounds in **v**_l_, and **A**^rev^ was its complementary block in **A**. To stay consistent with that convention in more general case, the columns corresponding to positive lower bounds need to be removed from **A**^irr^, and instead transferred to **A**^rev^.^3^ Thereby, **A**^irr^ corresponds to 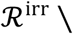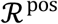 and **A**^rev^ corresponds to 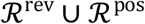. Whenever we speak of a factorization of the form Eq. (9) in the context of networks operating under general flux bounds, we also mean to incorporate this minor discretion.

The obtained factorization scheme demonstrates an interesting property of type-I nonstoichiometric BCs: the emergence of such BCs does not rely on the value of the flux bounds (specified by **v**_l_, **v**_*u*_). Consequently, BCs of this type originate from the network topology, and in particular, the existing irreversibility patterns. It follows that potential modeling errors in the form of inaccuracies in determining upper-and lower-bounds on fluxes has no impact on our ability to detect such BCs in a given network.

What is less desirable about them is the fact that they emerge in conjunction with blocked reactions in the network. Alternatively, suppose we first remove those blocked “bridge” reactions from the network. This would break down some linkage classes of the network into a number of smaller linkage classes, which means we would come up with a reduced network of slightly different linkage structure. This, in turn, expands the left nullspace of the incidence matrix **A**, that is, the matrix **U** will contain a higher number of columns, i.e. basis vectors. It follows that the stoichiometric factorization,

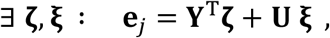

may have a larger solution set in the case of the reduced network. This means, the reduced network obtained by the removal of blocked reactions may accommodate a higher number of stoichiometric BCs. In fact, one can show that all type-I nonstoichiometric BCs of the original network appear as stoichiometric BCs in the reduced network.

A somewhat different phenomenon is the case of nonstoichiometric BCs that may only arise under non-canonical flux regimes, that is, when at least one irreversible reaction has a positive lower bound. We hereby refer to such nonstoichiometric BCs as <u>*type-II*</u>. As was shown in Section S1.4.1, all BCs must satisfy the implicit factorization given in Eq. (13). Assuming all blocked reactions have been removed from some network *G*, any remaining nonstoichiometric BCs in *G* must be of type-II. Clearly, the existence of type-II nonstoichiometric BCs in *G* relies on having a nonempty set 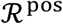.

Even though type-II nonstoichiometric BCs emerge in conjunction with the fixation of some reaction fluxes at upper- or lower bounds, as was shown in Section S1.4.2, they do not automatically rely on the network having some blocked reactions that effectively change its linkage structure. This is an interesting property of type-II nonstoichiometric BCs, because of which they can be seen as an unexpected nontrivial phenomenon in the steady state analysis,

Looking back at Eq. (13), the potential existence of a type-II nonstoichiometric BC encoded by **e**_*j*_ directly relies on the existence of feasible dual variables **λ**_u*t*_, **λ**_l*t*_, *t* = 1,2 that fit into the pattern **λ**_l*t*_ − **λ**_u*t*_ ∈ im(**A**^T^), *t* = 1,2. However, **λ**_l*t*_, **λ**_u*t*_, *t* = 1,2 are not free variables, but are tied to each other via other equations in Eq. (13), that is

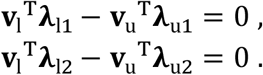

This casts light on an undesirable property of type-II nonstoichiometric BCs: the values **v**_l_, **v**_u_ appear explicitly in their factorizations. It follows that potential modeling errors in the form of inaccuracies in determining upper-and lower-bounds on fluxes may have a negative impact and result in a failure to correctly detect type-II nonstoichiometric BCs: if the value of some upper/lower bounds in the model (**v**_l_, **v**_*u*_) are modified or miscalculated, this could change the values of **λ**_l1_ − **λ**_u1_ and **λ**_l2_ − **λ**_u2_, which could means they would not necessarily satisfy the factorization for a given **e**_*j*_ anymore. This sensitivity of detection to the values of upper- and lower-bounds is an unwelcome feature with type-II nonstoichiometric BCs.

### 6 6S2 Analysis of biological networks

We categorized balanced complexes identified by Küken et al. [2] in twelve genome-scale metabolic models from organisms of all kingdoms of life, namely *A. niger* [3], *A. thaliana* [4], *reinhardtii* [5], *E. coli* [6], *M. acetivorans* [7], *M. barkeri* [8], *M. musculus* [9], *M. tuberculosis* [10], *N. pharaonis* [11], *P. putida* [12], *T. maritima* [13] and *S. cerevisiae* [14].

Balanced complexes were identified in networks including no blocked reactions; therefore, these networks do not include type-I nonstoichiometric BCs. To check if a BC *C*_*j*_ is a strictly stoichiometric BC we solve the feasibility problem in Eq. (18), where **e**_*j*_ is the vector with a unit value for the j^th^ entry and zero values elsewhere and **Y** being the stoichiometric map. The linear program will be feasible if **e**_*j*_ is a linear combination of rows in **Y**.

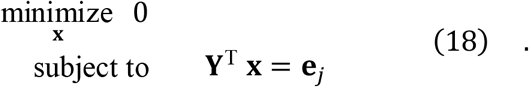

Moreover, Küken et al. identified BCs under different sets of constraints, including the scenario were reaction reversibility is considered and canonical flux bounds are imposed and the scenario were the lower bound of the biomass reaction is set to at least 99% of its optimum obtained from flux balance analysis [15]. Comparing the sets of BCs found under the respective scenarios allows to identify type-II nonstoichiometric BCs, which are those complexes being balanced under imposed lower bound on biomass, but not under canonical flux bounds. The remaining BCs are then classified as stoichiometric BCs.

## Footnotes

1 Otherwise, it can be shown that some irreversible reactions must be blocked.

2 We showed that 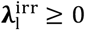 must be of the form 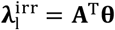. It follows that he algebraic sum of the entries in 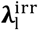 corresponding to any reaction cycle (loop in the graph) must be zero. Taking into account also the fact that 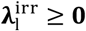and 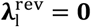, it follows that every entry of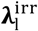 corresponding to a reaction inside a strong linkage class must be zero.

3 This replaces the original partitioning **A** = [**A**^rev^ **A**^irr^] into **A** = [**A**^zr^ **A**^nz^], where **A**^zr^ denotes the columns corresponding to (irreversible) reactions with zero lower bounds. by contrast to type-I nonstoichiometric BCs, which we associated with unsophisticated network reductions.

